# Cognitive resilience to Alzheimer’s disease in mice depends on gene-diet interactions

**DOI:** 10.1101/2025.02.07.637137

**Authors:** Amy R. Dunn, Harpreet Kaur, Yanchao Dai, Kevin Charland, Andrew R. Ouellette, Niran Hadad, Patricia H. Doyle, Glen H.G. Acosta, Elizabeth M. Litkowski, Timothy J. Hohman, Kristen M.S. O’Connell, Catherine C. Kaczorowski

**Affiliations:** The Jackson Laboratory, Mammalian Genetics, Bar Harbor, Maine, USA 04609; Washington University in St Louis, Department of Neuroscience, St Louis, Missouri, USA 63110; Tufts University School of Medicine, Boston, Massachusetts, USA 02111; University of Michigan Medical School, Department of Neurology, Ann Arbor, Michigan, USA 48109; University of Maine Graduate School of Biomedical Science and Engineering, Orono, Maine, USA 04469; Division of Bioinnovation and Genome Sciences, Translational Genomics Research Institute, Phoenix, Arizona, USA 85004; University of Kentucky College of Medicine, Department of Neuroscience, Lexington, Kentucky, USA 40506; Vanderbilt University School of Medicine, Department of Neurology, Nashville, Tennessee, USA 37232

## Abstract

Alzheimer’s disease (AD) has a complex etiology arising from largely unknown interactions between genetic and environmental (GxE) factors. Even in populations with causal familial Alzheimer’s disease mutations, there is variation in disease onset and progression, suggesting that clinical symptoms are modified by genetics and environment. Identification of such modifiers is critical, as mechanisms that promote resilience to high-risk AD mutations, unhealthy diet, or aging represent promising therapeutic targets for AD; global resilience factors that protect against multiple “hits” are among the highest priority for discovery. Both genetic and environmental protective factors in AD have been identified; however, GxE factors are incredibly difficult to study in human populations given complex genomes, poor self-reporting, limited data from underrepresented groups, and incompletely documented exposomes. Here, we (1) validate novel GxE tools using population of mouse strains that model the polygenic nature of human AD, (2) characterize individuals with cognitive resilience to high-risk genetic and dietary perturbations, (3) define and quantitate roles for genetics, sex, age, and diet, and (4) present data for the discovery of complex interactions that are nearly impossible to elucidate from humans or inbred mice. We found that a high-fat high-sugar (HFHS) diet is not universally damaging, as some strains showed an improved AD-related cognitive outcome when fed a HFHS diet, suggesting the need for personalized recommendations for dietary interventions in AD. Cognitive resilience to AD is polygenic; however, we found a locus on Chr 10 that was modestly associated with cognitive resilience to AD in females, and this association was strengthened by HFHS diet, pointing to an unexpected interaction between specific genetic loci and ‘unhealthy’ diet in AD risk and resilience. This study is the first of its kind to explore characteristics of AD resilience and GxE interactions in a genetically diverse mouse model. We present a subset of strains that exemplify global cognitive resilience to be leveraged for deep mechanistic studies aimed toward development of resilience-based, personalized therapeutic interventions.

**Highlights:**

- AD-BXD mouse models and data repository provide unprecedented tools for understanding individual differences arising from Gene x Environment factors in aging and AD.
- Evidence of global cognitive resilience to AD-relevant outcomes that is partially mediated by a resilience to aging and dietary risk factors.
- Cognitive resilience in AD-BXDs its polygenic nature, with a modest locus on Chr 10 whose association with resilience was amplified by a high-fat/high-sugar diet.
- The genetic factors mediating resilience to AD in humans also predict cognitive resilience in AD-BXDs, suggesting robust, translatable global resilience factors.

## 1.1 Introduction

Alzheimer’s disease (AD) is a common and irreversible neurodegenerative disease. It is also highly heterogeneous, with individuals displaying a wide range of symptom profiles, severity, progression, and pathology. Individual genetic variation explains some of this phenotypic variability [1]. Even among patients who harbor the same causative genetic mutations the disease onset, progression, and pathology are surprisingly variable [2–5], suggesting that there are additional genetic modifiers that influence the penetrance of these dominant mutations. Dozens of genetic variants have been identified that increase the risk of sporadic AD [6]. A handful of genetic variants have been shown to be protective against familial or sporadic AD [7–13] in addition to protective molecular, cellular, and environmental factors [14–16]. These protective, or resilience, factors have been increasingly recognized as representing particularly powerful therapeutic approaches to AD—rather than inhibiting “risk” pathways, we may be able to engage and leverage “protective” pathways to prevent disease progression in many individuals [11, 17–20]. Moreover, mechanisms of resilience and susceptibility may vary across individuals, with many molecular strategies leading to the same observed phenotype of resilience, or resilience in one or more cognitive domains vs others.

Despite all the known genetic contributors to disease susceptibility/resilience, we are still unable to account for a majority of predicted genetic risk for AD [1]. One hypothesis for this so-called missing heritability is that some genetic contributors interact with environmental factors, and these gene-environment (GxE) interactions may contribute significantly to AD risk and protection. These genetic contributors can exert different effects depending on environmental contexts: they may present small effect sizes in some environmental contexts or larger effect sizes when sensitized by other environmental factors. Conversely, known environmental contributors may only be meaningful depending on individuals’ genetic context. Diet, for example, is a commonly considered environmental factor that may modify risk of AD [21], with variations of diets high in fiber, low in sugar and saturated fat (e.g., a Mediterranean or MIND diet) currently being tested as a lifestyle intervention to prevent progression to (or of) dementia in older adults [21–24]. However, the reported effects of diet on cognitive function and AD risk are variable in both human and model organism studies [25]. The potential genetic factors contributing to the variable effects of environment are difficult to disentangle in human studies given the incredible complexity of the human exposome, and--with respect to diet--the incomplete record and control over diet over the human lifespan. Mouse models of AD, then, are a viable option for studying effects of environment/diet on AD and cognition given their highly controlled environments; however, mouse models of AD historically lack genetic diversity (or at least tractable genetic diversity), so true genome-wide GxE studies have been impossible.

Here, we use the AD-BXD model of AD, which integrates well-characterized causative AD mutations of the 5XFAD transgenic model of AD with tractable and replicable diversity of the BXD genetic reference panel [26–34]. This population provides an unprecedented opportunity to (1) identify resilient populations and explore genetic factors leading to resilience, and (2) experimentally probe genetic modifiers of effects of environmental risk and protective factors of AD and aging.

## 2. Methods

### 2.1. AD-BXD generation

AD-BXD mice were generated as described previously [26] by crossing female B6-congenic mice hemizygous for the 5XFAD transgene (MMRRC Strain #034848-JAX; RRID:MMRRC_034848-JAX) [28, 35, 36] with males from 38 BXD strains and parental B6 and D2 strains for 41 AD-BXD strains total [27] (Fig. 1A). The 5XFAD transgene includes five familial AD mutations total, three in the amyloid precursor protein gene (APP; Swedish mutation (K670N/M671L), Florida mutation (I716V), London mutation (V717I) and two in the presenilin 1 gene (PSEN1, M146L, L286V). We included both DBA/2J (JAX Strain #000671; RRID:IMSR_JAX:000671) and DBA/2J-Gpnmb^+^/SjJ (JAX Strain #007048; RRID:IMSR_JAX:007048) which has a corrected (functional) allele of *Gpnmb* as parental “D2” strains in this study, and B6D2F1 and B6xD2Gpnmb+ are abbreviated as “D2” and “D2G”, respectively. We included an average of 6 biological replicates per BXD strain, sex, 5XFAD status (nontransgenic or carrier), diet (chow or HFHS diet), and terminal age timepoint (3 mo, 6 mo, or 14 mo), for a total of 7,312 mice (4,239 chow-fed, 3,073 HFHS-fed) who entered the study and 6,164 mice (3,512 chow-fed, 2,652 HFHS-fed) who completed the study. Ns for each strain/sex/5XFAD genotype/diet/age group per phenotyping test are detailed in Table 1. Mice were group-housed in pens of 2-5 sex-matched littermates of mixed genotypes. To capture phenotypic and molecular features of “preclinical”, “early disease”, and “advanced disease” stages, we assigned mice to 3 mo, 6 mo, and 14 mo terminal timepoints, respectively. All mouse experiments took place at The Jackson Laboratory and were conducted in accordance with the JAX Institutional Care and Use Committee and the NIH’s Guide for the Care and Use of Laboratory Animals.

**Figure 1.**
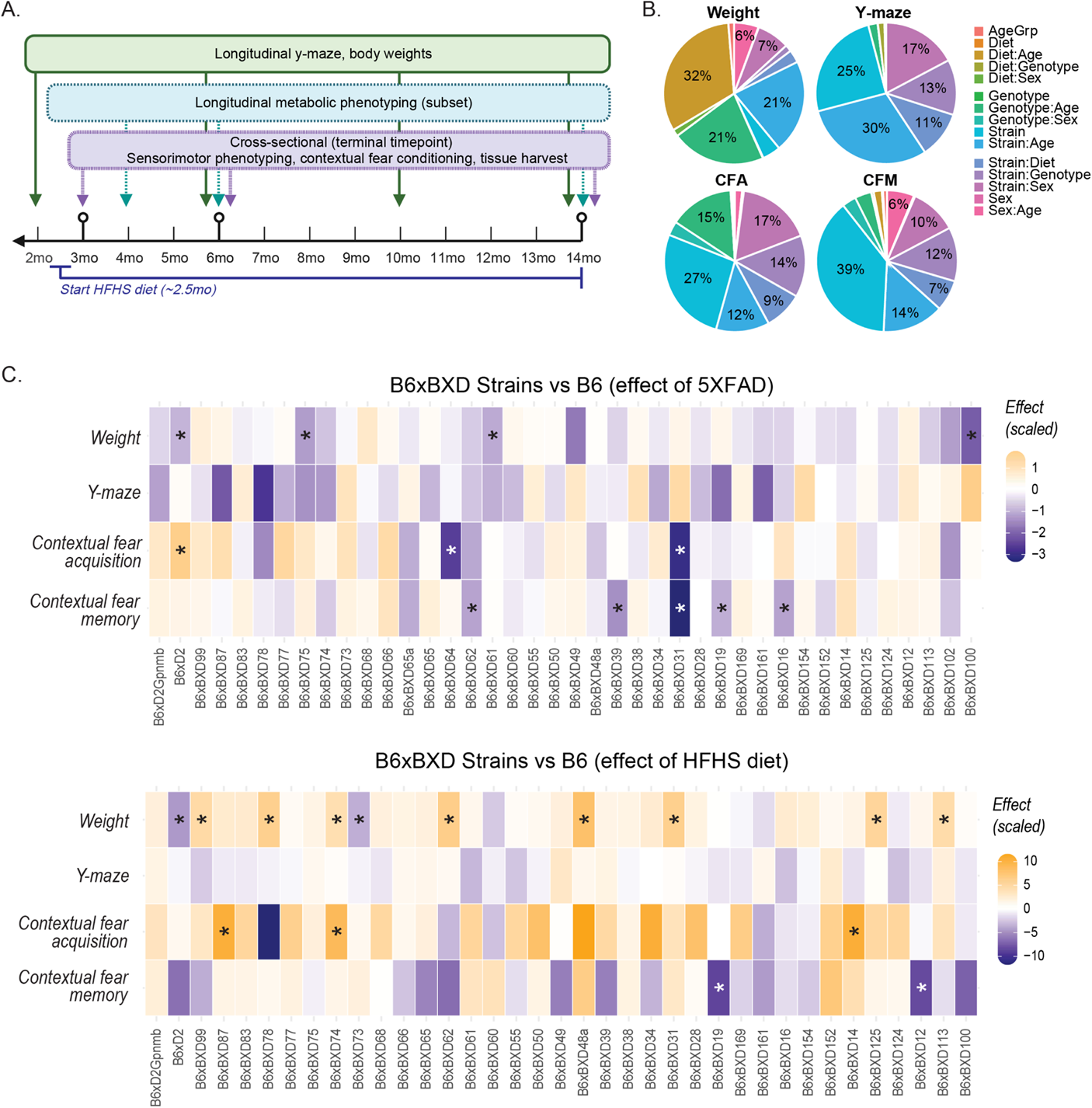
Study design and contribution of experimental factors on weight and cognitive outcomes. ***(A)*** Mice underwent baseline cognitive phenotyping at 2 mo and were placed on a HFHS diet at 2.5 mo or continued on a normal chow diet. They received cross sectional and longitudinal phenotyping through their terminal timepoints of 3, 6, or 14 mo of age. ***(B)*** Percent variance explained in weight and cognitive outcomes by all main variables and first-level interactions. ***(C)*** Visualization of linear mixed modeling results indicate that introducing genetic diversity to the 5XFAD model and to HFHS diet exposure leads to significant differences from B6, both in terms of the 5XFAD transgene and the effect of HFHS diet. On the color scale, orange = higher than B6; blue = lower than B6. Effect sizes have been scaled for visualization purposes, using the root square mean of the effect values. Asterisks indicate p < 0.05.

**Table 1.** Animal and strain numbers for each main phenotype, by sex, 5XFAD genotype, age, diet.

| Body weight: individual, (strain) n per group |  |  |  |  |  |
| --- | --- | --- | --- | --- | --- |
| Female |  |  |  | Male |  |
|  |  | 6 mo | 14 mo | 6 mo | 14 mo |
| Ntg | Chow | 584 (41 strains) | 360 (41 strains) | 487 (40 strains) | 320 (38 strains) |
|  | HFHS | 372 (38 strains) | 232 (38 strains) | 354 (37 strains) | 225 (36 strains) |
| 5XFAD | Chow | 579 (41 strains) | 360 (41 strains) | 547 (40 strains) | 326 (37 strains) |
|  | HFHS | 382 (38 strains) | 233 (38 strains) | 363 (36 strains) | 229 (37 strains) |
| Y-maze: individual, (strain) n per group |  |  |  |  |  |
| Female |  |  |  | Male |  |
|  |  | 6 mo | 14 mo | 6 mo | 14 mo |
| Ntg | Chow | 402 (41 strains) | 252 (41 strains) | 294 (40 strains) | 222 (38 strains) |
|  | HFHS | 303 (38 strains) | 210 (38 strains) | 286 (37 strains) | 207 (36 strains) |
| 5XFAD | Chow | 395 (41 strains) | 267 (41 strains) | 338 (40 strains) | 215 (37 strains) |
|  | HFHS | 320 (38 strains) | 218 (38 strains) | 287 (36 strains) | 202 (37 strains) |
| CFC: individual, (strain) n per group |  |  |  |  |  |
| Female |  |  |  | Male |  |
|  |  | 6 mo | 14 mo | 6 mo | 14 mo |
| Ntg | Chow | 304 (41 strains) | 305 (41 strains) | 264 (40 strains) | 236 (38 strains) |
|  | HFHS | 251 (38 strains) | 217 (38 strains) | 223 (37 strains) | 208 (36 strains) |
| 5XFAD | Chow | 307 (41 strains) | 309 (41 strains) | 306 (40 strains) | 234 (37 strains) |
|  | HFHS | 240 (38 strains) | 223 (38 strains) | 233 (36 strains) | 215 (37 strains) |

#### 2.1.2. Diets

Mice were fed ad libitum either normal control diet (“chow”; LabDiet 5K0G, 16% fat, 61% carbohydrates (0.69% sucrose) [37], 22% protein by kcal, or a high-fat/high-sugar (HFHS) diet; Research Diets D12451, 45% fat, 35% carbohydrate (17.5% sucrose), and 20% protein by kcal). Mice in the HFHS diet group were randomly assigned by pen to this diet condition and placed on HFHS diet at 2.5 mo of age following baseline cognitive testing at 2 mo of age.

### 2.2. Phenotyping pipeline

Mouse body weights were measured every two months, starting at 2 mo of age. Mice received longitudinal y-maze, every 2 or 4 mo throughout their lives starting at 2 mo of age, for a total of 12,639 individual y-maze tests. Body weights were taken at the same timepoints. A subset of mice (n=1-2 per strain/sex/5XFAD genotype/diet group) received comprehensive metabolic phenotyping longitudinally at 4, 6, and 14 mo of age, consisting of indirect calorimetry (IDC, 5 days), intraperitoneal glucose tolerance testing (IPGTT), and nuclear magnetic resonance (NMR) body composition analysis; of these metabolic phenotypes, NMR outcomes are included in the present report. At their terminal timepoints (3 mo, 6 mo, 14 mo), mice additionally underwent sensorimotor testing (grip strength, inclined screen, and narrow beam tests) and contextual fear conditioning (CFC), which measured short-term memory (contextual fear acquisition, CFA, as measured by percent freezing in the post-shock 4 period) and long-term memory recall (contextual fear memory, CFM; see Fig. 1A). Before each behavioral testing timepoint, mice were habituated to transport and the testing room for one hour each day for three days. Each apparatus was thoroughly cleaned with 70% ethanol before and after tests to minimize detectable olfactory cues produced by previously tested mice. Details of each phenotype are as follows.

Y-maze: To measure spatial working memory, we conducted y-maze tests as described previously [38, 39]. Mice were placed in one arm of a three-arm plexiglass maze and allowed to explore for seven minutes. Visual cues were placed outside of each arm. Movement through the maze was tracked with AnyMaze software (Stoelting Co, Wood Dale, IL, USA). Entries into each of the three arms before returning to a previously-entered arm were recorded as “spontaneous alternations.” Percent correct spontaneous alternations was calculated as [spontaneous alternations/(total arm entries – 2)]*100. Mice who entered fewer than three arms during the testing period were excluded from analysis.

Sensorimotor: Sensorimotor function was assessed using narrow beam (balance and motor coordination), inclined screen (geotaxis), and grip strength (muscle strength) as described previously [32]. During the narrow beam test, mice were placed on a raised beam (1.2cm width) with a dark box and an open platform at either end. Latency to reach the end of the beam was recorded; faster times indicate better motor function. Animals who fell off the beam or who did not complete the task received the maximum score of 300 s. During the inclined screen assay, mice were placed head-down on a grid raised at a 45-degree angle. Latency to rotate to head-up orientation was recorded. Animals who fell off the grid or who did not complete the task received the maximum score of 300 s. During the grip strength test, mice were placed with their forearms gripped to a grid and pulled backward. Force in newtons required to pull the mouse off the grid was recorded as forelimb grip strength. A sensorimotor composite score was calculated by averaging the z-score-transformed score (r scale() function, which calculates the root square mean, [x - mean(x)) / sd(x)]), on each component (narrow beam, geotaxis, grip strength).

Nuclear magnetic resonance (NMR) analysis of body composition: To measure lean mass, fat mass, and water mass, animals underwent NMR body composition analysis. Awake animals were enclosed in a Plexiglas tube and placed into an Echo MRI (Houston TX, USA), where body composition was measured according to manufacturer’s protocols.

Indirect calorimetry: Mice were placed individually in Promethion metabolic phenotyping cages (Sable Systems International) for five days. Mice were allowed to acclimatize to the cages for one light period. Gas exchange, food and water intake, and activity were averaged over 12hr light, 12hr dark, and 24hr periods for the remainder of the 5-day testing period. Respiratory quotient was normalized to metabolically-adjusted weight (lean body mass + (0.2 fat mass) [40].

Contextual fear conditioning (CFC): To measure contextual fear acquisition and memory recall, we conducted CFC. Mice were placed in fear conditioning chambers (Actimetrics, Wilmette, IL, USA) and allowed to explore for a baseline period of 150 s. They received four mild foot shocks (0.9 mA, 1 s duration) with an intershock period of 140s +/- 10 s [38, 41–43]. Each “post-shock” period was defined as the 40 s following the shock. Movement and percent time spent freezing were recorded using FreezeFrame software (Actimetrics). Percent time spent freezing in the fourth postshock period (PS4) was recorded as an index of short-term CFA (memory acquisition).

Twenty-four hours later, mice were returned to the chambers with the same context and allowed to explore for 10 min without receiving foot shocks. Percent time spent freezing during the 10 min exploration was recorded as long-term memory recall (contextual fear memory, CFM). Animals who exhibited >50% freezing during the baseline acclimation period prior to training were excluded from analysis. CFC was administered once per animal immediately before euthanasia and tissue harvest.

### 2.3. Survival analysis

While this study did not include a formal longevity paradigm, we did assess successful survival to the latest timepoint (14 mo). We categorized early deaths as “health-related” or “censored” study removals. “Health-related” deaths included spontaneous death due to age or illness, euthanasia for veterinary health reasons, and death due to aggressive behavior (aggressor or target). “Censored” deaths were not due to the health/behavioral status of the mouse and included euthanasia required as the result of environmental conditions, e.g., inability to group-house due to death of cage-mates. We then calculated rate of survival (excluding censored deaths) to 14 mo within each strain/5XFAD genotype/sex/diet group as a measure of survivability and attrition.

### 2.4. Enzyme-linked immunosorbent assay (ELISA)

Following the completion of the phenotyping paradigm, mice were euthanized using isoflurane and brains were removed and hypothalamus, hippocampus, midbrain, cerebellum, and isocortex (frontal and rear) were sub-dissected for subsequent biochemical analysis. A total of 422 samples were collected from 41 strains n=395 5XFAD and n=27 non-transgenic mice (for non-pathological controls) at three different ages (3 mo, n= 137 (66M/71F), 6 mo, n=144 (67M/77F), 14 mo, n=141 (65M/76F) of both diets were used for quantification of beta-amyloid 1-42 (Aβ1-42). ELISAs were performed as described previously [26, 28]. Snap-frozen cortical tissues (post frontal lobe removal; ∼25mg) were homogenized in PBS containing 1% TritonX-100 + protease inhibitor cocktail (Thermo Scientific, 78429) + DX reagent (Qiagen, 1011008) with a 5mm stainless steel bead (Qiagen, 69989) using a TissueLyser II system (QIAGEN) at 30s^-1^ for 3 min. Homogenates were then sonicated for 2×10 seconds and protein concentration was measure using NanoDrop 2000 UV-Vis Spectrophotometer (ThermoScientific, USA). Sample were diluted to 10µg/µl and mixed with 5M Guanidine HCl (GuHCl) for the final amyloid beta extraction. The GuHCl extraction was performed overnight at 4℃ on orbital shaker. Samples were then diluted appropriately and run in duplicate along with standards using a commercially available ELISA kit (Cat# 298-62401, FUJIFILM Wako Chemicals, VA, U.S.A). The absorbance was read at 450 nm using a microplate reader (SpectraMax® i3x) from Molecular Devices LLC., CA, USA. The levels of Aβ1-42 are reported as picomoles per liter (pmol/L).

### 2.5. QTL mapping

Genome-wide QTL mapping was performed as described previously [39] with the R package R/qtl2 [44] using a Haley-Knott regression method and adjusting for kinship with the leave-one-chromosome-out (LOCO) method. Additive covariates were determined by identifying significant main effects of factors (Sex, Diet, 5XFAD Genotype, Age) on each given trait. Significance levels were determined using the scanone() function in R/qtl2 using 1,000 permutations.

### 2.6. Statistics

Statistics and data visualization were performed in R. Significant effects of diet/sex/age/AD status on cognitive and noncognitive outcomes were identified using ANOVA, linear regression, or linear mixed modeling as described in results. Multiple comparisons were corrected for with a Bonferroni adjustment and post-hoc comparisons from ANOVA were achieved with a Tukey’s HSD test where appropriate and as noted.

#### Calculation of a polygenic resilience score

We classified strains in a binary approach as being “susceptible” or “resilient” based on composite resilience scores at 14 mo: chow-fed females with higher than median resilience scores were classified as “resilient” and lower-than-median scores were classified as “susceptible”. We next selected seventeen genes that have previously been associated with human AD resilience in Seto, et al. 2021[17] and Lopera, et al. 2023 [7], and identified the marker SNPs representing alleles (homozygous for the B6 allele, i.e. BB, “B”; or heterozygous for the B6 and D2 alleles, i.e. BD, “D”) at those genes across the BXD strains. We next calculated the number of susceptible and resilient strains that have a B or D allele at each genetic locus to generate a ratio of susceptible/resilient strains that harbor a B or D allele at each locus. We then classified either the B or D allele at each locus as conferring resilience to generate an allele-specific “resilience score”. We log-transformed the allele resilience score and summed these resilience scores across alleles for each strain to generate strain-specific polygenic resilience scores.

## 3. Results

We generated a population of 41 AD-BXD strains and their nontransgenic (Ntg-BXD) littermates for deep behavioral characterization before and during HFHS diet (Fig. 1A). We selected the early adulthood timepoint (6 mo) as a reference timepoint for baseline cognitive and metabolic function, and the middle aged timepoint (14 mo) to capture an advanced disease stage for subsequent analyses. The number of individuals and strains represented in each age/sex/genotype/diet group is shown in Table 1. Numbers per group varied slightly due to study attrition over time, strain availability over time, and animal husbandry/experimental requirements preventing singly housed animals or single-genotype pens.

To assess the effect of age, diet, 5XFAD genotype, BXD strain, and sex on metabolic and cognitive outcomes across individuals, we performed a type III ANOVA on phenotype measures (Fig. 1B). Age, sex, and background strain contributed significantly to body weight (Age: 1.3% of the variance, p=1.03e-18; Sex: 0.1% of the variance, p=0.01; Strain: 4.2% of the variance, p=4.37e-33), albeit to a lesser extent than variance explained in body weight by interacting factors. Diet * Age interactions explained the most variance in body weight (32%, p=0), followed by 5XFAD Genotype * Age (21%, p=2.71e-278) and Strain * Age (21%, p=3.61e-239). Y-maze working memory was significantly influenced by Strain (25%, p=0.002), and again interactions between experimental factors contributed significantly to the variance in working memory (Strain * Age: 30%, p=6.58e-05; 5XFAD Genotype * Age: 2%, p=0.01; Diet * 5XFAD Genotype: 1.5%, p=0.04). For short-term memory, once again Strain was the only factor with a significant main effect, accounting for 27% of the variance in contextual fear acquisition (p=3.06e-12), and interactions between factors contributed to a majority of the remaining known variance (Strain * Sex: 17%, p=4.14e-05; 5XFAD Genotype * Age: 15%, p=5.74e-18; Strain * 5XFAD Genotype: 14%, p=0.002; Strain * Age: 12%, p=0.02). For long-term memory, Strain and Age contributed to phenotype variance (Strain: 39%, p=2.48e-45; Age: 1%, p=0.007) and interacting factors contributed a significantly to the remaining known variance (Strain * Age, 14%, p=1.94e-45; Strain * 5XFAD Genotype: 12%, p-1.28e-07; Strain * Sex: 11%, p=1.45e-05; Strain * Diet: 7%, p=0.01). Supplemental Table 1 details the full Type III ANOVA models. The relatively large contribution of interacting factors to each of these traits suggests that these variables interact to influence metabolic and cognitive phenotypes in a strain-specific manner. We also calculated the heritability [45] of these traits and found, consistent with our ANOVA results, that metabolic and cognitive traits were generally modestly heritable ([G_var_/(G_var_+E_var_)], where G_var_ = variance across strains and E_var_ = average within-strain variance); average overall h^2^ range = 0.07-0.57; Supplemental Table 2), again indicating that genetics contributes to a proportion of these traits, but that additional factors contribute to disease-related outcomes.

### Introducing genetic diversity to the 5XFAD model results in significantly varied cognitive and metabolic effects associated with AD and HFHS diet

Next, we leveraged linear mixed modeling with strain as a random factor to compare all strains to the reference strain, B6. First, we assessed the interaction between strain and 5XFAD status (Phenotype ∼ 5XFAD*Strain + (1 | Strain)). Figure 1C, top panel, shows the z-scaled effect size of the 5XFAD transgene on each strain across body weight and cognitive outcomes. Several strains showed AD-related outcomes that significantly differed from B6 in body weight and contextual fear conditioning, denoted by an asterisk (p < 0.05). Likewise, HFHS had significantly different effects on body weight and contextual fear conditioning outcomes on many strains compared to B6. These analyses highlight the role of interacting genetics factors in regulating response to AD-causative mutations and environmental sensitizers like a HFHS diet.

To further assess the impact of GxE interactions on the progression and severity of AD-related phenotypes, we evaluated outcomes in 3 mo and 14 mo old Ntg and AD-BXD mice fed either chow or HFHS diets. Consistent with previous BXD and AD-BXD reports, there was a broad population-wide variation in disease-related outcomes such as body weight, cognitive performance and survival in both diet groups (Supplemental Fig. 1)[26, 27, 32, 46, 47]. At a population level, HFHS diet did not result in significantly negative impacts on cognitive function at 3 or 14 mo of age, nor did it result in a global increase in mortality by 14 mo of age. However, to assess the specific strain differences in response to HFHS diet in males and females of each AD genotype, we fit a linear mixed model to evaluate the interaction of Diet and Strain with Strain as a random factor in 14 mo old mice (Weight ∼ Diet*Strain + (1 | Strain)) to account for baseline strain-to-strain variability in body weight. In each sex/5XFAD genotype group, over half of strains (28/40 ntg female strains, 21/40 AD females, 24/37 ntg males, 24/37 AD males) did not show a significant effect of diet on body weight. To identify strains with significant effects of diet on cognitive traits, we again used linear mixed models as described above. HFHS diet did not significantly impact y-maze performance in the majority of strains (number of strains where y-maze was affected by HFHS diet was 0/41 strains in AD or Ntg females, 2/38 strains in Ntg males, 0/37 strains in AD males); however, a few strains did show a significant effect of diet, both positive and negative, on short-term but not long-term memory outcomes (CFA: 2/41 Ntg female strains, 5/41 in AD female strains, 0/38 Ntg male strains, 8/37 male strains; CFM: 0/41 Ntg or AD female strains, 0/38 Ntg male strains, 0/37 AD male strains). Full linear mixed model outcomes are detailed in the supplemental data (Supplemental Tables 3-18).

We also observed significant variation in cognitive trajectory over time in mice regardless of diet. In general, we observed AD-related cognitive decline in female mice across all cognitive domains; HFHS did not exacerbate cognitive decline in any domain in mice on a C57BL/6J background (Supplemental Fig. 2A) or in the AD-BXD population as a whole (Supplemental Fig. 2B). Specifically, in females, we observed a significant effect of both 5XFAD genotype and age by ANOVA followed by a Tukey’s HSD (multiple comparisons of means) test, where 5XFAD females showed worse working memory performance (effect of 5XFAD transgene was −1.63 percentage points, p=0.006). Female mice also showed decline in working memory starting at 6 mo of age (2-6 mo, p-adj=0.003; 2-10 mo, p-adj = 4.01E-05; 2-14 mo, p-adj = 6.58E-05), and 5XFAD females exhibited a decline in short- and long-term memory (CFA: 3-14 mo, p-adj<1e-07; CFM: 3-14 mo, p-adj=0.0001). Male mice exhibited decline in working memory only by 10 mo of age (2-10mo, p-adj=0.003; 2-14mo, p<1.0E-07), and HFHS-fed males showed better working memory performance overall (p-adj=0.008). This observation that AD-related cognitive impairment occurs earlier and is more pronounced in females reflects the phenomenon in humans and animal models that females are more susceptible to AD [48–52].

### A subset of strains show robust, global resilience against age- and AD-related cognitive decline

Resilience to AD is challenging to study in human populations, because of limited information regarding cognitive abilities throughout life, challenges accessing brain tissue from presymptomatic ages, and a lack of information regarding diet and other environmental exposures. Thus, characterizing resilience in a genetically diverse mouse population such as the AD-BXDs provides an unprecedented opportunity for discovering various molecular, genetic, and behavioral strategies that may all present as cognitive resilience.

To capture global cognitive resilience in AD, we incorporated all cognitive domains assayed (Y-maze/working memory, contextual fear acquisition/short-term memory, and contextual fear memory/long-term memory) and calculated a global resilience score. We took advantage of the reproducibility of the AD-BXD strains to adapt a common measure of human resilience—residual variance from the ‘expected’ cognitive function—to mice. This approach is used across human studies, where expected performance may be modeled given known characteristics (genetic risk, age, education, amyloid load, etc.) of the participants [53, 54]. In our AD-BXD population, rather than modeling expected performance, we instead can consider the healthy, non-aged adult (i.e., 6 mo) performance of Ntg, chow-fed animals as the “expected” performance based on that strain’s genetic background. Then, how much each strain deviates from this expected performance with age, transgene status, or environmental ‘hits’ (e.g., HFHS diet) is the residual variance that may be used to quantify relative resilience/susceptibility of that strain to these factors. This is visualized in Supplemental Fig. 3, where the line of slope = 1 is a theoretical expected performance if strains performed identically at 14 mo to their Ntg/6mo/chow-fed baseline (i.e., if they experienced no age-, diet-, or AD-related cognitive decline). We repeated this for each cognitive domain (working, short-term and long-term memory), and with a composite cognitive score incorporating all three assays (Fig. 2A). The composite cognitive score was calculated by scaling each test with a z-transformation (r scale() function, which calculates [x - mean(x)) / sd(x)]), and summing the z-scores within each strain/sex/5XFAD genotype/diet group. We measured distance from “expected” performance for each strain/sex/diet/5XFAD genotype/age group for each cognitive domain. Finally, these values were scaled with a z-transformation and summed to obtain a “composite residuals” score. In Fig. 2B, residual performance for each Ntg and AD-BXD strain at 14 mo is quantified and plotted in order from least-to-most resilient female, chow-fed AD strain. Composite residuals scores ∼0 or higher indicate that a strain performed similarly at 14 mo to their 6 mo/Ntg/chow-fed counterparts and represent cognitive resilience. We used this measure as a quantitative, continuous trait for further analyses of resilience and susceptibility. To estimate the contribution of genetics to this resilience score (i.e., heritability), we fit a linear model with BXD strain as the predictor variable. This revealed an R^2^ of 0.40, suggesting that genetic background predicts about 40% of the variance in resilience (p = 0.004).

**Figure 2.**
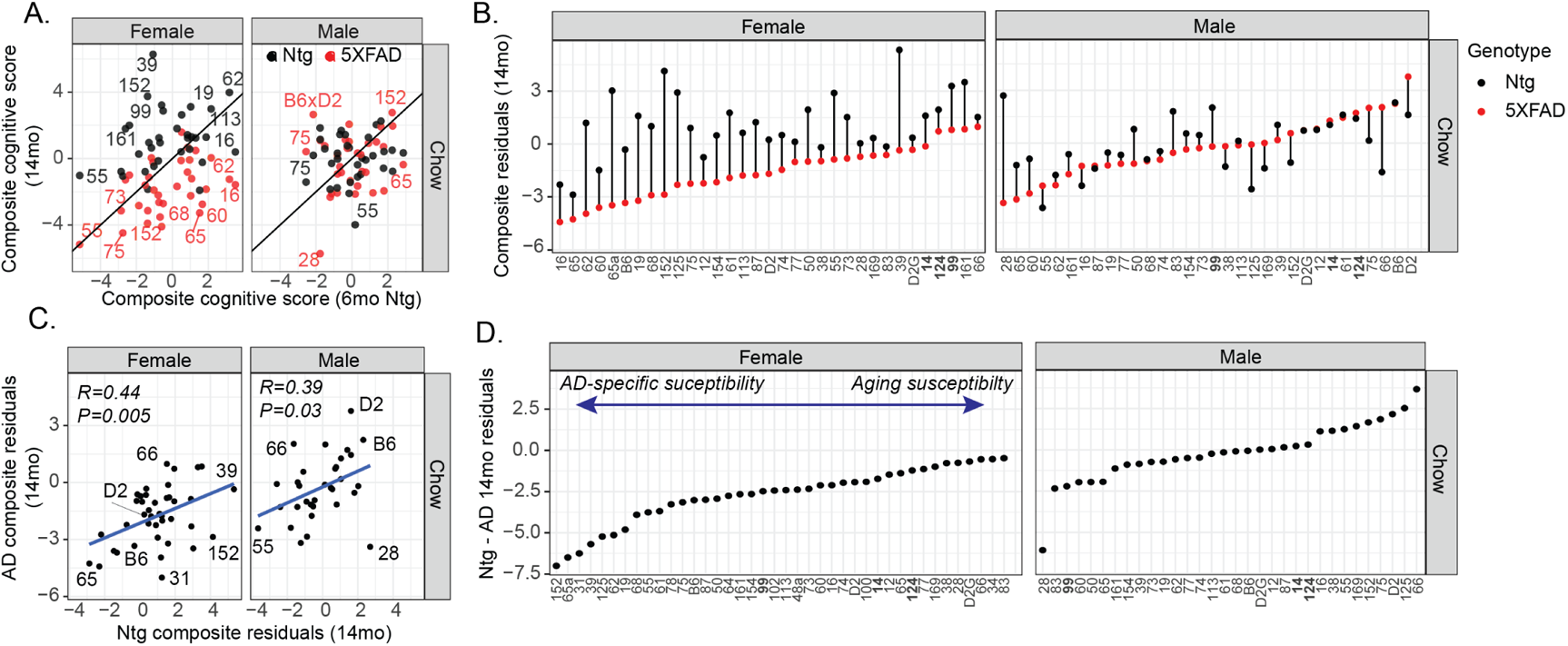
Cognitive resilience to Alzheimer’s disease mutations varies across AD-BXDs and is partially mediated by normal cognitive aging. ***(A)*** We calculated a composite cognitive score at 6 mo and 14 mo by summing the z-scored performance on three cognitive tests (y-maze, contextual fear acquisition, and contextual fear memory. Here, we plot 14 mo performance by strain-matched 6 mo Ntg baseline performance. Black line at *y = x* shows theoretical performance if strains performed exactly at at 14 mo compared to 6 mo Ntg strain-matched controls. Strains that are above this line perform relatively better than 6 mo Ntg controls; strains that fall below this line perform relatively worse compared to 6 mo Ntg controls. ***(B)*** We calculated residual variance as distance from “expected” (6 mo Ntg control) performance for each cognitive domain in 14 mo Ntg and AD-BXDs as a measure of resilience/susceptibility to cognitive decline. All female AD-BXDs, but not all males, showed greater susceptibility to cognitive decline compared to strain-matched Ntg controls. ***(C)*** To estimate the relationship between susceptibility to normal cognitive aging and AD-related cognitive decline, we plotted composite cognitive residuals in AD-BXDs against their strain-matched Ntg controls. There is a significantly positive relationship in both females (R=0.44, p=0.005) and males (R=0.39, p=0.03), with an adjusted r2 of 0.30 when controlling for sex. Thus, resilience to normal aging accounts for about 30% of resilience to AD-related cognitive decline. ***(D)*** We then plotted strains by the difference between their Ntg and AD cognitive residuals. Strains with more negative scores showed greater decline when expressing the 5XFAD transgene compared to normal aging (i.e., AD-specific susceptibility); strains with positive or less-negative scores had comparable or less decline when expressing the 5XFAD transgene compared to normal aging (i.e., similar susceptibility/resilience in AD and normal aging).

Next, we calculated the difference in composite resilience scores between Ntg and AD-BXDs to assess the relative contribution of a general *resilience/susceptibility to cognitive aging* in observed resilience and susceptibility to AD-sensitized cognitive decline (Fig. 2C-D). Normal aging accounted for a significant amount of the variance in AD cognitive decline, as evidenced by an overall adjusted R^2^ of 0.3 (p < 2.25E-09) and individual within-group correlations between 0.39-0.45 (Fig. 2C). Thus, much of an individual’s resilience to AD may be mediated by a more general resilience to aging. However, there were exemplary strains that showed AD-specific vulnerability or resilience. In females, mice from strain 152 were relatively *resilient* to age-related cognitive decline, while showing significant *susceptibility* to AD-related cognitive decline. In chow-fed females, strain 66 showed especially resilient phenotypes in both AD and Ntg mice.

### Resilience to AD-related cognitive decline is mediated by gene-environment interaction

Next, we asked whether resilience factors are similar in chow-fed vs HFHS-diet fed animals. First, we assessed the relative overall contribution of HFHS to AD-related cognitive susceptibility/resilience and found no interaction between Diet*5XFAD genotype on our composite residual score in females (p = 0.36), indicating that HFHS neither exacerbated nor protected against AD-related cognitive decline in females. Using linear modeling, we found that the composite residual score in chow-fed animals accounts for about 40% of the variance in their HFHS-fed counterparts (r^2^ = 0.39; within-group male and female correlations shown in Fig. 3A), indicating that there are drivers of resilience that may act similarly regardless of diet. In line with this, some strains consistently scored higher on cognitive resilience regardless of sex, AD mutation status, or diet. Our linear model results (Supplemental Tables 3-18) also showed that one strain, B6xBXD124 scored significantly higher on resilience overall (estimate = +1.85, p=0.033). Two other strains showed a statistical trend toward global resilience, B6xBXD14 (estimate = +1.12, p=0.197), and B6xBXD99 (estimate = +1.32, p=0.127). Trajectories of these strains’ cognitive function between 3-14 mo of age, together with mice on a B6 background for contrast, are visualized in Supplemental Fig. 4. This observed cognitive resilience does not appear to be related to survival bias: cognitive resilience is not mediated by a strain’s 14 mo survival rate in a multiple linear regression model with 5XFAD genotype, sex, diet, and survival rate as predictors (df=1, F=2.2; p = 0.14 for survival rate). The degree of resilience in these strains hint at robust, cross-domain and environment-independent resilience factors that may be genetically controlled.

**Figure 3.**
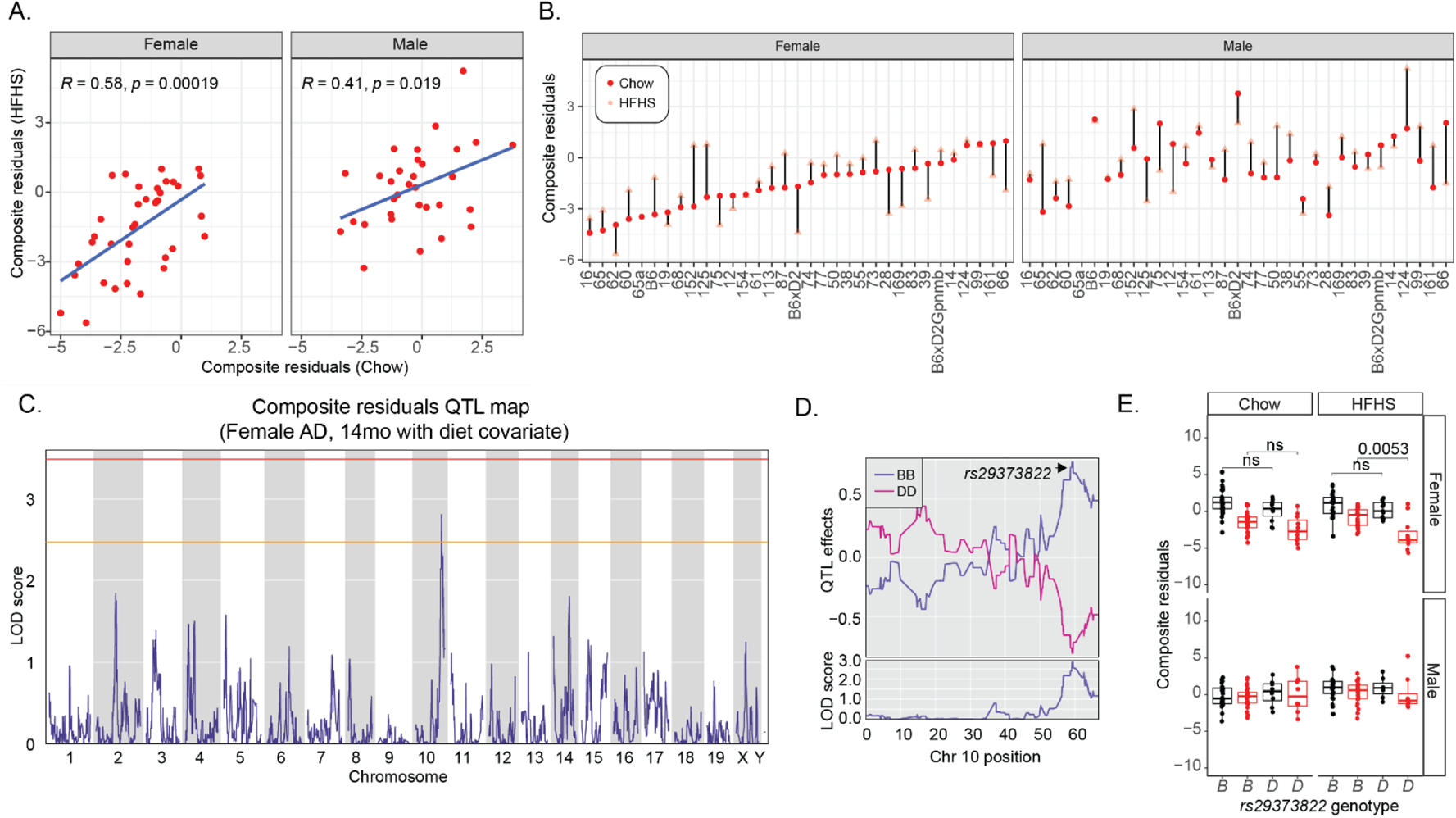
Diet modifies cognitive resilience in AD. ***(A)*** Comparing composite cognitive residuals in HFHS diet fed AD-BXDs compared to strain0matched chow-fed controls shows a strong correlation between cognitive resilience regardless of diet in females (R=0.58, p=0.00019) and males (R=0.41, p=0.019). An adjusted r^2^ of 0.37 indicates that resilience in chow-fed animals accounts for about 37% of the variance in resilience in their HFHS-fed counterparts. This is also represented in ***(B)***, where strains are ordered from least-to-most resilient when fed normal control chow. Several strains show greater or lower resilience depending on diet. ***(C)*** We performed quantitative trait locus (QTL) mapping in 14 mo AD females with diet as an additive covariate. We found a suggestive peak on Chr. 10 that was modestly associated with cognitive resilience. ***(D-E)*** The peak marker rs29373822 was associated with poorer cognitive function in females fed a HFHS diet only in individuals carrying the *D* allele at this locus.

On the other hand, several strains show wide variation in resilience depending on sex, diet, or AD mutation status. For example, female B6-5XFAD mice are highly susceptible to cognitive decline (composite resilience score ≈ −3.5) when fed a chow diet, and this susceptibility is partially attenuated when fed a HFHS diet (composite resilience score ≈ −1). Males of the same strain, B6-5XFAD, are resilient to cognitive decline regardless of diet (composite resilience score ≈ +2.5). Such cognitive resilience to the 5XFAD transgene in males on a B6 background is consistent with our previous findings [38] and highlights the important sex differences in AD-related phenotypes.

### Resilience to cognitive decline in AD females is polygenic, and this genetic association may be modified by diet

Next, using this global resilience score as a quantitative trait, we performed quantitative trait locus (QTL) mapping to determine if cognitive resilience in advanced AD is strongly modified by one or more loci, and if any genetic association is modified by diet. Likely reflecting the polygenic nature of resilience, we did not find a single locus that was significantly associated with global cognitive resilience; however, mapping resilience in 14 mo old female mice with diet as an additive covariate revealed a suggestive QTL on Chr 10 (Fig. 3C). The *B* allele at this locus was associated with stronger cognitive resilience (Fig. 3D-E) in HFHS-fed females. This locus was only associated with resilience in females (p-adj = 0.042; Fig. 3G), supporting data from us and others that the genetic architecture of resilience varies by sex [8].

The suggestive locus on Chr 10 (defined as the interval around the peak remaining within 1.5LOD of the peak) is a 10.07MB region, Chr10:117.30-127.37MB with the peak at 121.64MB. The locus contains 176 genes, including 72 named genes and 105 gene models and unannotated genes. The peak marker SNP is *rs29373822*. This D allele at this SNP was significantly associated with lower cognitive resilience in 5XFAD females fed a HFHS diet (nominal p = 0.0053; p-adj = 0.042; Fig. 3E), but not in females fed a chow diet or in males. This SNP was not associated with differences in noncognitive outcomes (e.g., body weight, sensorimotor function, Aβ1-42; data not shown) in any group.

### Human resilience genes predict resilience in female, but not male, AD-BXD mice

Following this, we probed the translatable value of our model by assessing contribution of genes known to be associated with human resilience to global resilience in AD-BXDs. We identified variants across the BXD genomes in loci containing known human AD resilience genes, with human resilience defined by residual cognitive variance quantification analogous to our present quantification [17]; human resilience genes without variants between B6 and D2 genomes were excluded from this analysis. First, we calculated a “polygenic resilience score” (PResS) [38, 55] using homologs of loci previously associated with resilience in humans, including *App, Apoe, Picalm, Tmem106b, Casp7, Abca1, Abca7, Sorl1, Rab10, Rest, Plcg2,Treml2, Dlgap2, Klotho, Bdnf, Ms4a6c*, and *Reln [7, 17]*. The PResS was correlated with resilience in AD-BXD females regardless of diet, but not in males (Fig. 4A), supporting previous findings that genetic mechanisms of resilience vary by sex [8]. In a model with sex and diet covariates, the PResS still significantly predicts resilience, with an adjusted r^2^ of 0.19 (p = 8.87e-07). Importantly, this indicates that the genetic factors governing resilience in humans are relevant in AD-BXDs, underscoring their value as a tool to understand the mechanisms of resilience to AD and cognitive aging. Three observations suggest that the genetic drivers of resilience may vary by environmental context: first, the PResS was more strongly associated with resilience in chow-fed females vs. HFHS-fed females. Second, three individual loci (*Abca1, Sorl1, Tmem106b)* were significantly associated with cognitive resilience at a nominal level in chow-fed females (D allele associated with better resilience), but not in HFHS-fed females. The D allele at the *Dlgap2* locus was significantly negatively associated with resilience in females regardless of diet (Fig. 4B).

**Figure 4.**
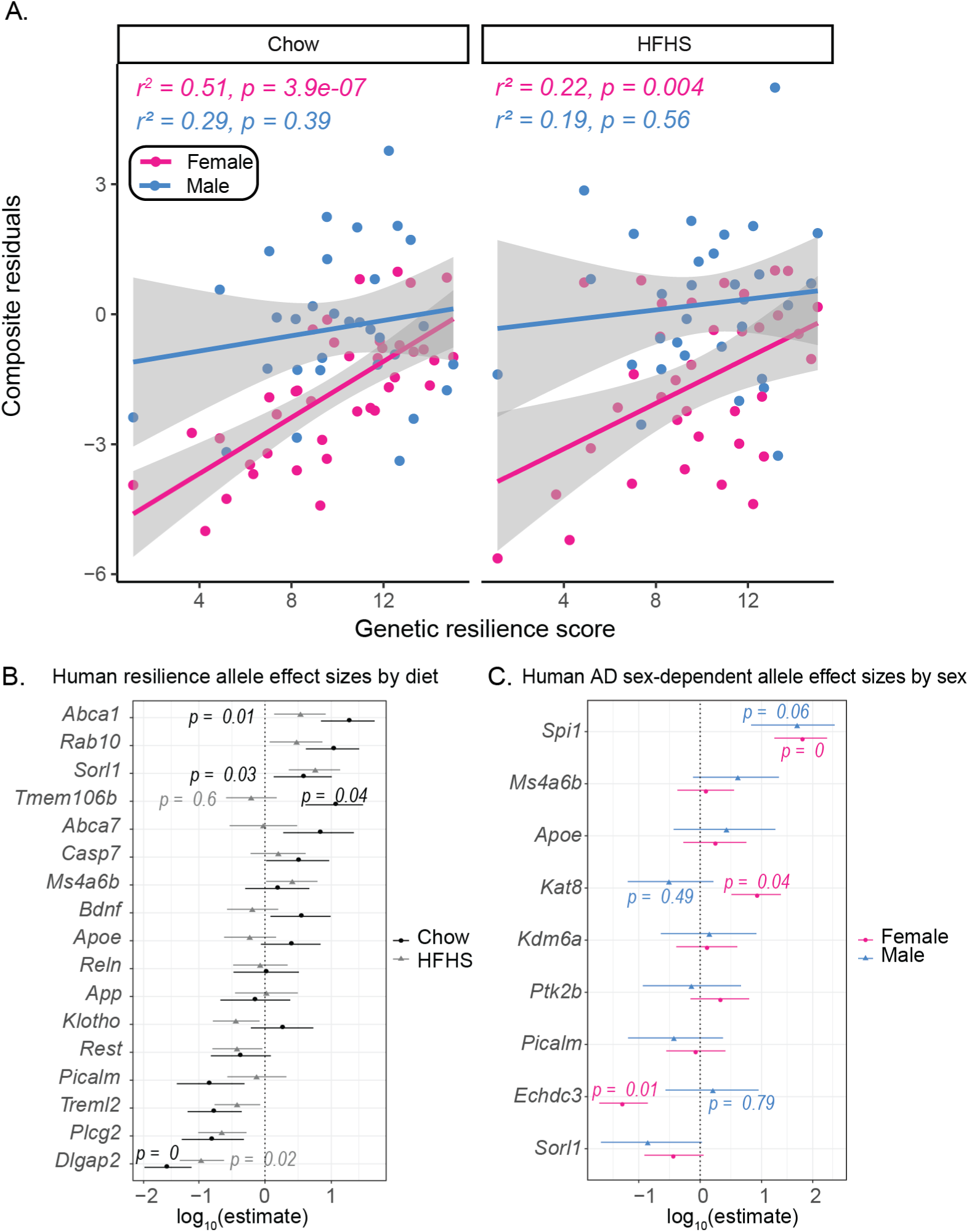
Human AD cognitive resilience genes predict cognitive resilience in AD-BXDs. ***(A)*** We calculated a polygenic resilience score (PResS) from 17 genes previously associated with resilience in human AD. This PResS predicted cognitive resilience in AD-BXDs in a sex-specific manner, with a stronger association in females (chow: r^2^=0.51, p=3.9e-07; HFHS: r^2^=0.22, p=0.004) compared to males (chow: r^2^=0.29, p=0.39; HFHS: r^2^=0.19, p=0.56) in either diet condition. ***(B)*** To evaluate the relative contributions of marker variants for each of the 17 genes, we used a linear model to calculate odds ratios, plotted here on a log scale. Several of these genes individually predicted resilience in chow-fed, HFHS-fed, or both, in females. In chow-fed animals, the D allele of *Abca1*, *Sorl1*, and *Tmem106b* were associated with higher resilience but not in HFHS-fed animals. The D allele of *Dlgap2* was associated with worse cognitive outcomes in both chow- and HFHS-fed females. ***(C)*** Finally, given the sex differences in AD-related cognitive decline, we assessed the relative association of several previously-identified sex-specific AD resilience genes. We found three genes that were significantly associated with resilience in females but not in males (D allele associated with higher resilience in females only, *Spi1* and *Kat8*; D allele associated with lower cognitive resilience in females only, *Echdc3*).

Finally, we asked whether genetic drivers of cognitive resilience are related to metabolic or other non-cognitive factors. At a nominal significance level, the PResS was associate with a handful of metabolic traits in HFHS-fed females only, particularly basal metabolic rate as quantified by respiratory quotient (see Methods, Supplementary figure 5; R=0.41,p=0.017), fat mass (R=-0.4, p=0.02), and glucose tolerance (R=-0.41, p=0.017). However, formal mediation analyses with the mediation.test() function from the r “bda” package did not indicate a significant mediating effect between these traits and the relationship between the PResS and cognitive resilience in females fed a HFHS diet (GTT AUC: Sobel z-value=0.07, p=0.946; body weight: Sobel z-value=-1.08, p=0.28; respiratory quotient: Sobel z-value=0.27, p=0.79), suggesting that AD resilience genes influence multiple traits independently.

Next, to specifically test whether human sex-dependent genetic modifiers of AD risk impact cognitive outcomes in AD-BXDs, we assessed variants proximal to genes reported to have differential effects on AD risk in males and females [8]. Of this list, three genes (*Spi1, Kat8, Echdc3*) showed female-specific associations with our global cognitive resilience score. In humans, *SPI1* and *ECHDC3* variants are specifically associated with female resilience to AD, while *KAT8* variants are associated specifically with male resilience to AD (Fig. 4C)[8].

## 4. Discussion

The AD-BXDs are a uniquely valuable tool that has allowed us to assess resilience and susceptibility to AD. Moreover, they have provided the first opportunity to evaluate gene-environment interactions in AD in a genome-wide manner, revealing that resilience and GxE contributors to resilience are polygenic traits.

### Susceptibility and resilience to cognitive decline are heterogeneous traits not predicted by individual demographic factors

In AD-BXDs, the degree of cognitive decline by 14 mo of age was highly variable across strain, sex, and diet. We did not observe a robust or global effect of HFHS diet on cognitive trajectory, but rather a dependence on the genetic background of the strain across the panel. HFHS diet was either neutral, protective, or damaging in terms of cognitive resilience in AD. Our finding that several strains differed significantly from B6 in terms of effect of HFHS diet on body weight gain and cognitive function highlights the value of introducing genetic diversity to understand disease mechanisms and risk factors. Though often underappreciated, there is considerable variability in the reported effects of a HFHS diet on AD-related outcomes[56–60]. As in our study, HFHS diets have been found to be damaging, neutral, or protective; this previously reported variability may have depended on experimental feeding paradigm (e.g., acute vs chronic HFHS diet), age, model, genetic background, sex, dietary makeup, and outcome measure. In fact, starting HFHS diet early in life, prior to AD pathology onset, is protective in some mouse models of AD [57], which may also reflect (1) the particularly damaging effects of inconsistent or “yo-yo” dieting or weight cycling throughout adulthood [61] and/or (2), the “obesity paradox” where excessive body weight in mid and late life has been found to be neuroprotective [62]. Interestingly, in studies that did find HFHS exacerbation of AD-related outcomes, these effects appear to be independent of the metabolic effects of HFHS [63], which is also consistent with our finding that the variability in cognitive outcomes was not related to variability in metabolic outcomes in our AD-BXD population. Given the variability across the literature in AD diet study designs, it has been difficult to draw conclusions about contributing factors that lead to varied outcomes. In our present results, the variability we found was striking, but our study design using a genetic reference panel--the AD-BXDs--provided an unprecedented opportunity to study both the strain differences in response to HFHS diet on AD-related cognitive phenotypes, and the differences in predictors of cognitive trajectory in chow- vs HFHS-fed animals.

Our study does provide additional support to a significant body of evidence that females are more susceptible to AD than males [48, 51, 52, 64]. Consistent with human disease, we observed the greatest susceptibility to AD-related cognitive decline in female AD-BXD mice. In fact, in all chow-fed female strains, AD mice exhibited greater cognitive decline compared to their Ntg counterparts. Male sex and HFHS diet resulted in improved cognitive outcomes in several strains, when compared to female or chow-fed genetic counterparts.

We hypothesized that some degree of resilience to AD was due to underlying resilience to general aging processes. In fact, we found a significant association between resilience to “normal” cognitive decline in middle aged Ntg animals and resilience to AD-related cognitive outcomes. These data suggest that there are likely common mechanisms between successful normal cognitive aging and cognitive resilience to AD, adding to the literature showing common genetic and mechanistic bases of cognitive resilience/decline in normal aging and disease [65, 66]. However, many strains did exhibit selective vulnerability to AD. For example, in chow-fed females, strain B6xBXD152 is highly resilient to cognitive decline by age 14 mo in Ntg animals, but highly *susceptible* to cognitive decline when harboring the 5XFAD transgene. Thus, some individuals likely harbor genetic susceptibility factors that overwhelm other factors that would otherwise promote successful cognitive aging.

To add to this complexity, some strains show dramatically different cognitive trajectories based on sex. To use B6xBXD152 again as an example, B6xB6152 females are particularly susceptible to AD-related cognitive decline; however, B6xBXD152 males are highly resilient to AD-related cognitive decline. BXD strains such as these may be particularly valuable in studying sex-dependent susceptibility and resilience in AD.

### Genetic predictors of resilience do not generalize to males in AD-BXDs

Several genes have been identified in previous studies as promoting cognitive resilience in humans [8, 17]. Using a polygenic resilience score (PResS) generated using variants across these genes in the AD-BXDs, we found that these human resilience factors are also associated with resilience in females, but not strongly in males. The PResS was significantly associated with cognitive resilience in chow-fed AD females, regardless of metabolic outcomes. However, a portion of the protective effect of these resilience drivers was mediated via their association with glucose tolerance in HFHS-fed AD females. In HFHS-fed AD females, the PResS was associated with better glucose tolerance (lower AUC on an IPGTT), suggesting that these genetic drivers of resilience may also improve metabolic outcomes on HFHS, which, in turn, lends protection against cognitive decline in AD. Thus, maintenance of metabolic health may be important in maintaining cognitive health, particularly in HFHS-fed individuals. We also examined known sex-specific modifiers of AD, and we found that a few of these modifiers (*SPI1, KAT8, and ECHDC3*) were uniquely associated with cognitive resilience in female AD-BXD mice but not in males. Finally, when we performed QTL mapping to identify novel mediators of cognitive resilience in AD-BXDs, our peak locus on Chr10 was associated with cognitive resilience in females, but not in males. Previous literature has suggested that the genetic architecture of resilience is sex-dependent, and our data support these findings. Our observation that the association between variants in these genes and resilience may also be modestly modified by diet, also hints that while genetic factors may be robust across environmental factors, the degree of protection may be influenced by environmental context.

Our suggestive resilience locus on Chr10 contains several promising candidate genes that could promote resilience in AD. One gene within this locus, *Slc16a7, w*hich encodes the protein neuronal monocarboxylate transporter MCT2, was previously identified by our group as mediating CFM in genetically diverse mice under various dietary interventions (e.g., caloric restriction)[67]. This strengthens our confidence that this locus may be one locus of many that is important in regulating dietary effects on cognitive function in aging and AD. Additionally, the several annotated genes contain missense mutations and thus may also represent candidates that are plausibly involved in the complex phenotype of cognitive resilience: *Cdk4*, kinase that is involved in response and metabolism of lipids and has been associated with obesity and type 2 diabetes; and *BC048403*, which encodes a subunit of KICSTOR, a component of the mTORC signaling pathway targeted by known resilience-promoting therapies such as rapamycin [68, 69]. Further validation of these candidates is warranted to evaluate the robustness of these candidates, as they may be potentially leveraged to promote resilience in people at risk for AD.

### GxE in AD resilience: HFHS may be detrimental or protective, depending on genetic context

Previous studies have investigated how single AD-related genes may interact with certain environmental risk or protective factors that impact AD risk [70–84]. For example, educational attainment and lifetime cognitive activity are most strongly protective in *APOE4* carriers [80, 85], and *APOE4* carriers are particularly susceptible to AD risk associated with obesity, low physical activity, heavy metal exposure, and sleep disturbances[71, 75–77] [see [74] for a review of known GxE interactions in AD]. Thus, we expected some variability in response to HFHS in the AD-BXDs, and we sought to characterize these gene-by-diet interactions in a population-wide manner.

We identified several strains whose susceptibility and resilience varied by diet, underscoring the individual variability in response to environmental (and likely treatment) interventions. This is important to consider for future pharmacotherapeutics, but also for commonly recommended lifestyle interventions, such as the WW-FINGER studies [86–90] that show efficacy in preventing cognitive decline. For example, we observed several strains that would likely be characterized as good candidates for a low-fat, low-sugar dietary intervention, such as B6xBXD12 AD females, that were susceptible to cognitive decline on a HFHS diet but not on a low-fat diet. Conversely, our data suggest that some individuals may not benefit—and indeed may experience exacerbated cognitive decline--from low-fat, low-sugar diets: strain B6xBXD125 females, as an example, showed significant improvement of AD cognitive outcomes when fed a HFHS diet vs a low-fat, low-sugar diet.

One goal of these analyses was to identify models of the most robust examples of cognitive resilience in AD. Strains that showed resilient cognitive outcomes despite aging, high-risk AD mutations, and diet, may be considered “super resilient” strains: these strains were resilient to cognitive decline in aging and across cognitive domains, even with AD mutations and a HFHS diet. B6xBXD124, 99, and 14 were some such strains and thus may be good candidates for further studying robust mechanisms of resilience against genetic and environmental “hits” in AD.

It is worth noting that we and others have previously characterized B6 as a resilient strain and the B6 genome as harboring resilience factors [26, 91]. Our current study differs from these previous studies by scale (∼900 mice in our previous study versus ∼7,000 in the current study) and power to detect sex differences. In fact, we did observe resilience in male B6 mice; only female B6 mice were susceptible to AD. Furthermore, female B6-5XFAD mice had improved cognitive outcomes when fed a HFHS diet. This sex difference highlights not only the need to study both sexes in AD and in AD resilience in sufficient numbers, but it also indicates that genetic and environmental contributors to resilience are different between males and females. Moreover, these data underscore the limited translational value of using only B6 males in preclinical interventions testing—a strategy which, heretofore, has been largely unsuccessful in effective drug development.

### Translatability & Conclusion

This study represents a significant step toward understanding resilience to AD and in understanding individual differences in susceptibility to genetic and environmental risk factors. We have identified several strains that will be ideal candidates to help elucidate the nature of resilience to AD, whether that is global cognitive resilience, sex differences in resilience, or resilience to environmental sensitizers of AD. We have also demonstrated that integrating multiple cognitive domains to capture global cognitive function results in a high degree of translatability to the human disease, where our “global residuals” scores showed genetic alignment with the human disease and AD resilience; this underscores the superior translatability of AD-BXDs to the human disease and provides a resource for deeper mechanistic studies using strains that are exemplars of resilience and susceptibility to AD.

## Acknowledgements

The authors would like to thank Timothy Leach, David Anderson, Grace Fisher, Natalia Bachelder Quattrochi, Sophie Martin, Kathleen Vollmer, Gregory Richard, Catherine Witmeyer, and Amanda Adams-Jewett for their contributions to mouse husbandry and phenotyping and Brianna Gurdon for critical review of manuscript drafts. Additionally, the authors gratefully acknowledge the expert assistance of several JAX Scientific Services with the work described in this publication: Drs. Vivek Philip and Jigang Zhang and Computational Sciences at The Jackson Laboratory for data management support, the Center for Biometric Analysis at The Jackson Laboratory for metabolic phenotyping support, and the Transgenic Genotyping Service at The Jackson Laboratory for transgenic mouse genotyping support. The Animal Care and Clinical Laboratory Animal Medicine staff at The Jackson Laboratory provided excellent care for each mouse that was a part of this study.

This work was supported by NIH RF1AG059778 (KMSO and CCK), Alzheimer’s Association AARF-18-565506 (ARD), Alzheimer’s Association AARF-22-928454 (HK), NIH R01AG054180 (CCK), NIH R01AG057914 (CCK)

## Author contributions

Conceptualization: ARD, HK, TJH, KMSO, CCK

Conducted experiments: ARD, HK, KC, ARO, PHD, GHA

Data analysis: ARD, HK, YD, KC, NH

Writing, figures: ARD, HK

Writing input: PHD, MD, NH, EML, TJH, KMSO, CCK

Supervised study: KMSO, CCK

Review and approval of the final manuscript: All authors

## Data availability statement

All experimental data have been submitted to the mouse phenome data repositories, Mouse Phenome Database (phenome.jax.org) and GeneNetwork (genenetwork.org) and will be freely accessible with no restrictions before publication (accession code and public access pending curator review; also available at https://github.com/KMSOConnellLab/ADBXD_GxE). BXD genotypes are available at GeneNetwork (gn1.genenetwork.org/genotypes/BXD.geno)

## Supplemental figures

**Supplemental figure 1.**
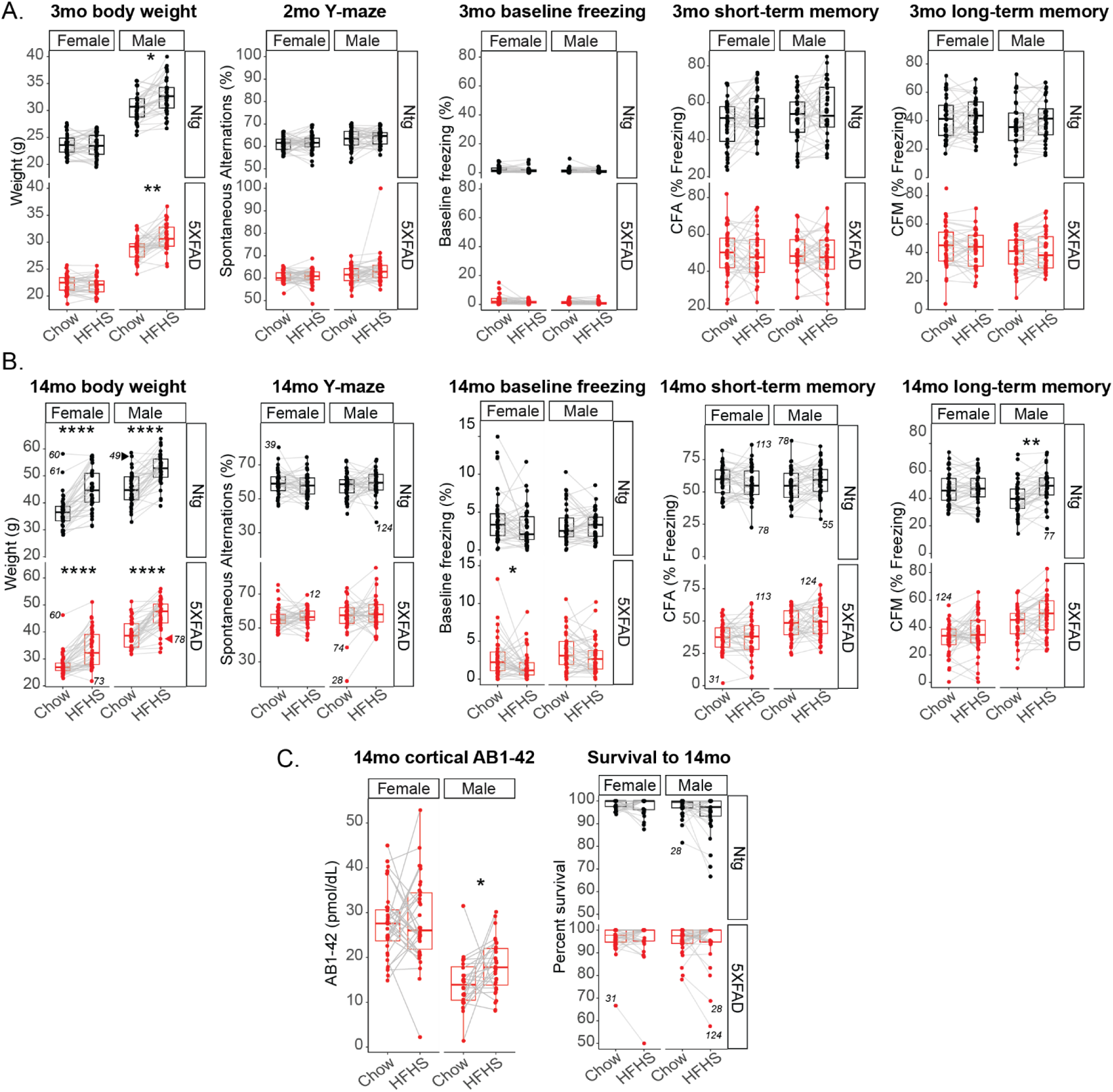
Population-level effect of HFHS at 3 mo and 14 mo of age. Individual strains have variability in direction and degree of body weight and cognitive response to HFHS diet at (A) 3 mo and (B) 14 mo. (C) At 14 mo, females fed a HFHS diet had similar levels of cortical abeta1-42; males fed a HFHS diet had slightly higher abeta1-42 at 14 mo. HFHS diet had no effect in any group on survival to 14 mo.

**Supplemental figure 2.**
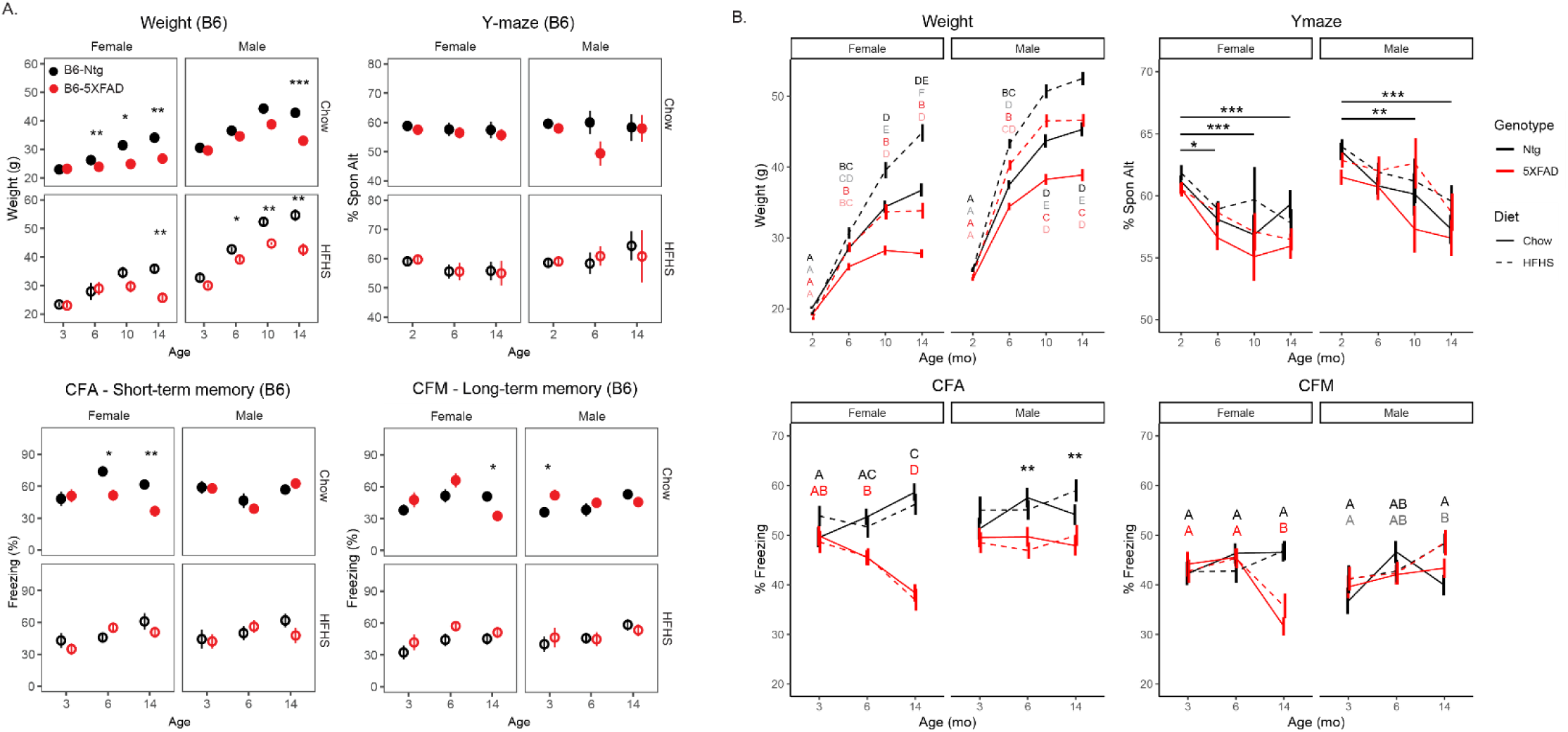
Trajectories of weight and cognitive function in mice on a homozygous C57BL/6J background and Ntg- and AD-BXDs by sex, genotype, and diet. (A) Body weight and cognitive function trajectories in B6 mice with and without the 5XFAD transgene, on normal control chow and HFHS diet. B6 mice with the 5XFAD transgene exhibited AD-related metabolic and cognitive symptoms, with significantly reduced body weight compared to Ntg controls starting at 6-14 mo of age, depending on sex/diet. Only females on a chow diet showed impaired cogtnive function relative to Ntg controls, with significantly reduced short-term memory by 6 mo and significantly reduced long-term memory by 14 mo. (B) Across the AD-BXD population, body weight varied significantly by age, genotype, diet, and itneractions between these vacotrs in both males and females by ANOVA. Differing letters denote statistically significant differences between diet/age/genotype groups. By ANOVA, y-maze performance varied by age (females) and diet, with no interaction between these factors. Asterisks denote statistically sifnicnat difference in performance across age groups. In females, short-term memory (CFA) varied by age, genotype, and an interaction between these factors. Differences between age/genotype groups are denoted by differing letters. In males, short-term memory (CFA) varied by genotype; asterisks denote significant genotype differences in short-term memory at 6 mo and 14 mo. Finally, long-term memory (CFM) varied by age and genotype (females) or age and diet (males). Significant differences between age/genotype (red/black) or age/diet (solid/gray) groups are denoted by differing letters.

**Supplemental figure 3.**
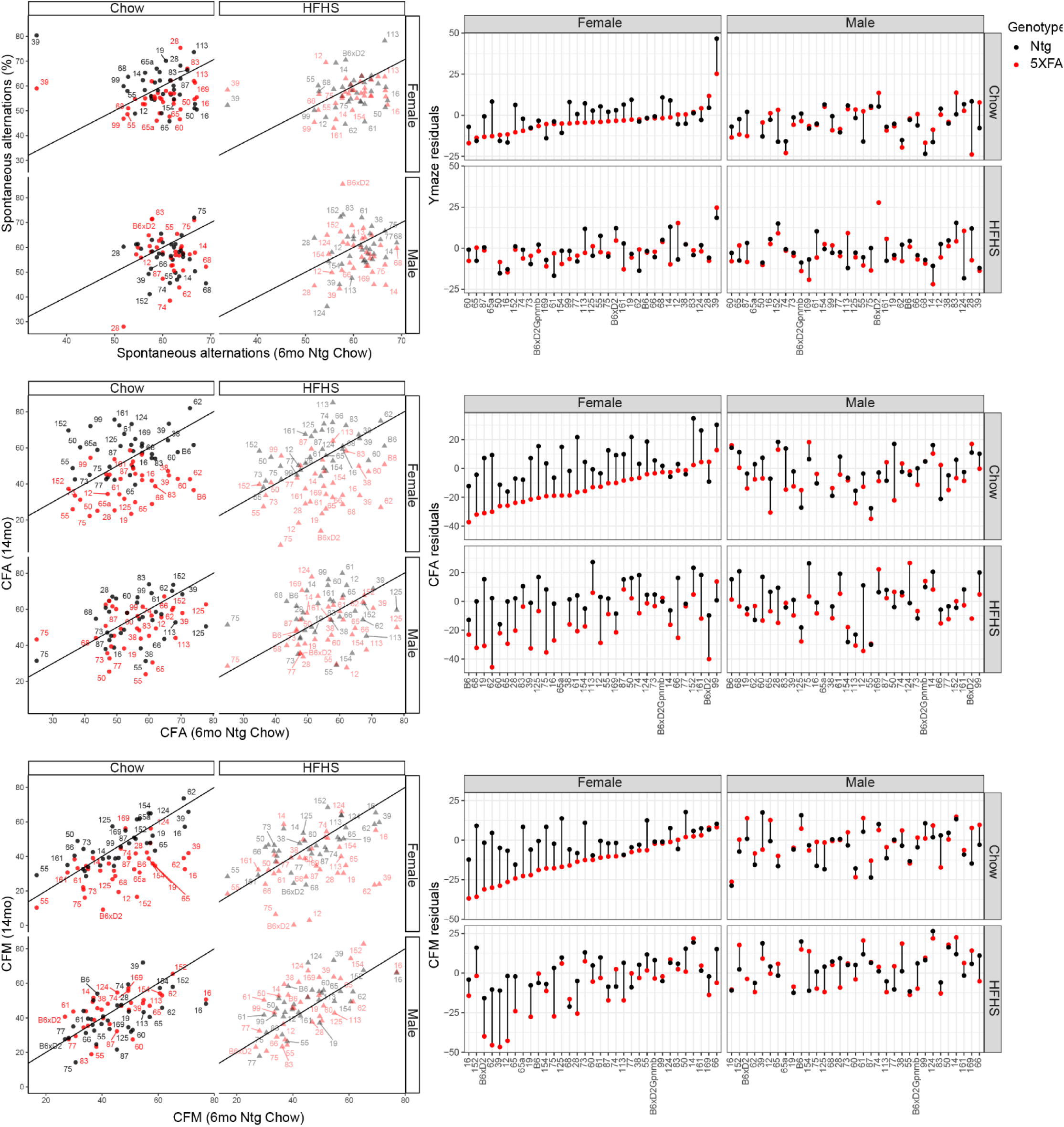
Calculating residual variance from “expected” baseline performance for each cognitive assay that contributed to the “composite” residual score. (Related to figures 2-3)

**Supplemental figure 4.**
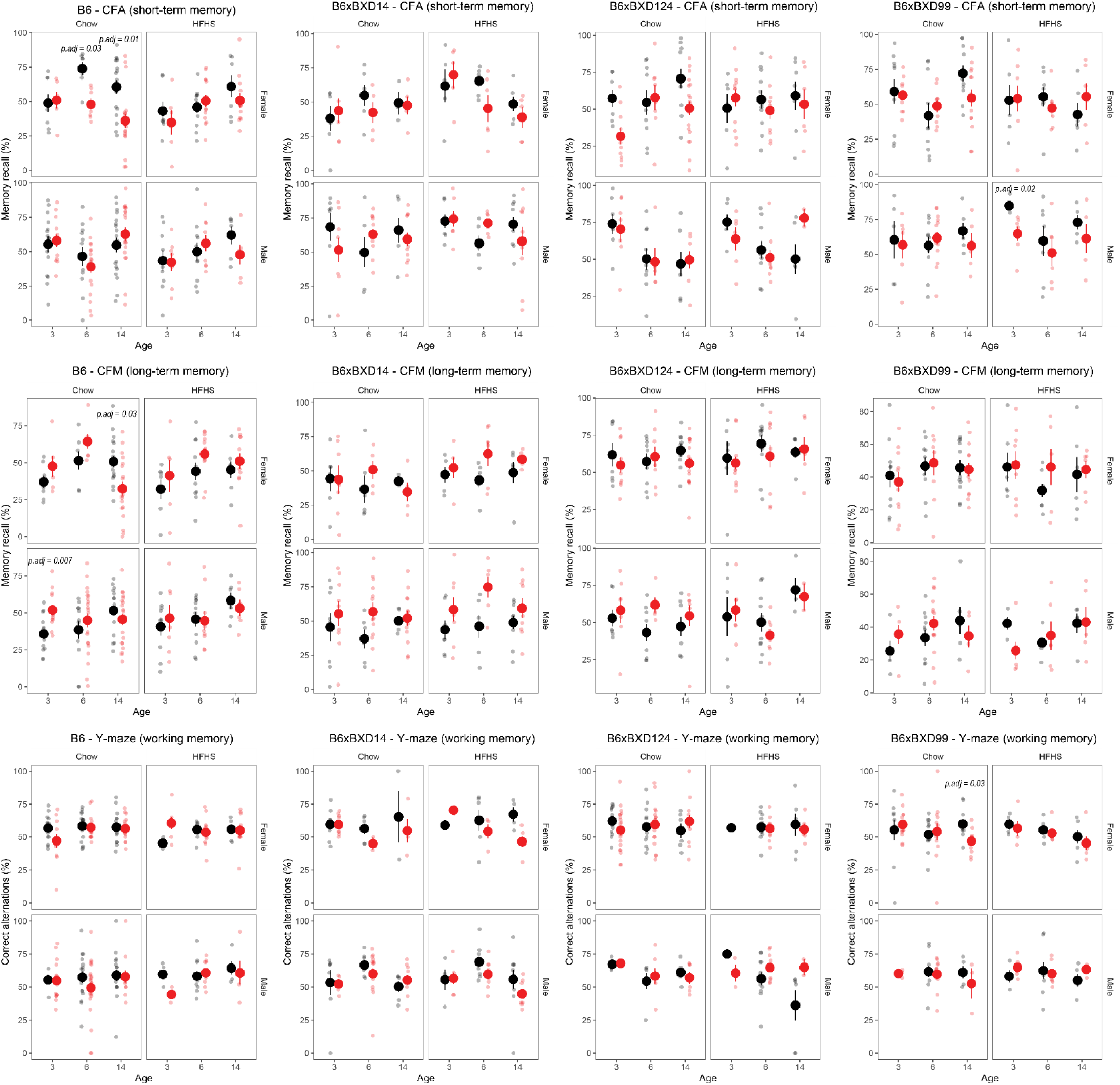
Individual strain trajectories for working memory (y-maze), short-term memory (CFA), and long-term memory (CFM). These strains were selected to highlight exceptional resilience to decline, and shown in contrast to B6 mice—females of which are vulnerable to cognitive decline as measured by CFC traits.

**Supplemental figure 5.**
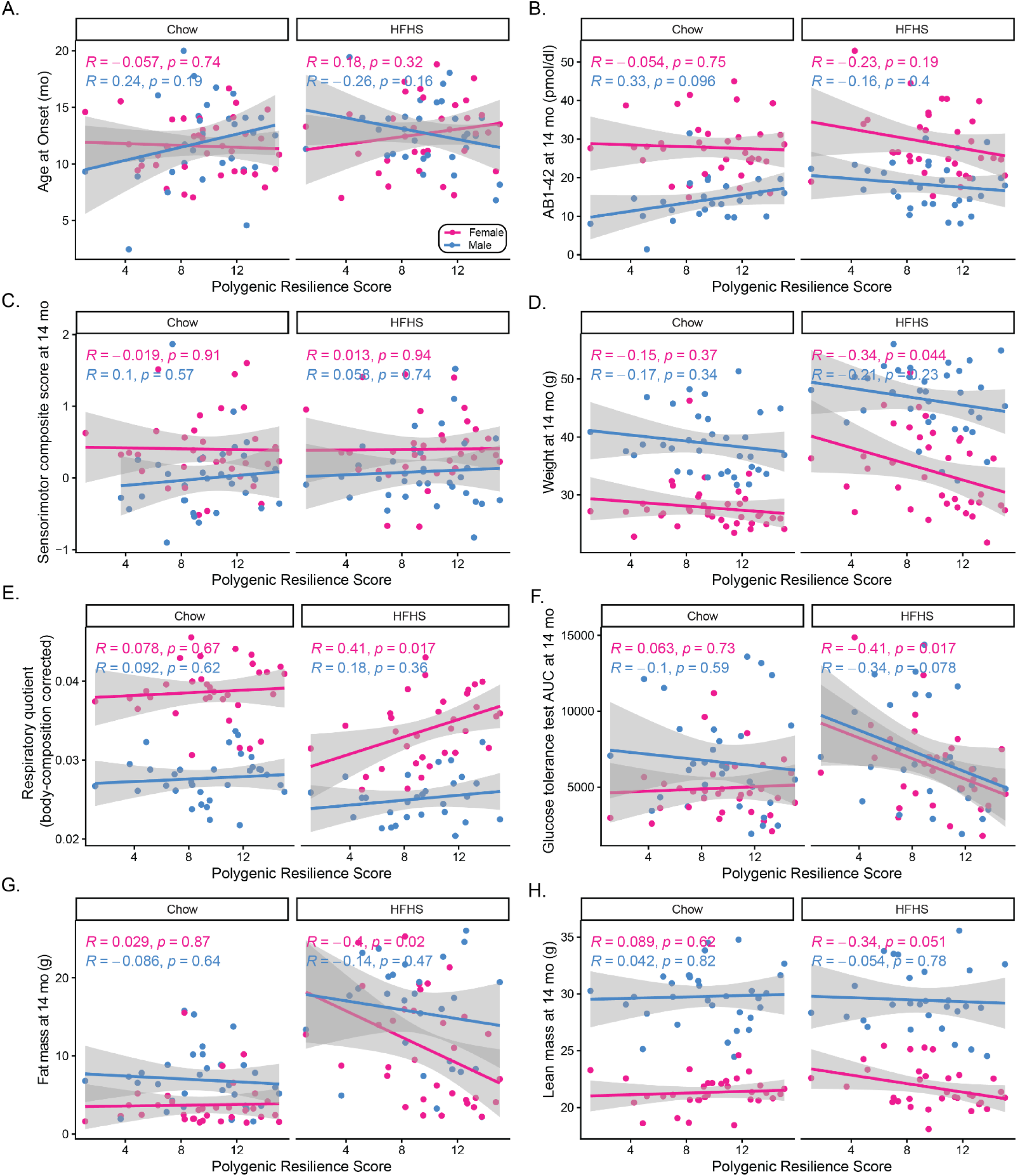
Correlations between polygenic resilience score and noncognitive traits in the AD-BXDs.

